# Predation, evo-devo, and historical contingency: A nematode predator drives evolution of aggregative multicellularity

**DOI:** 10.1101/2025.05.04.652091

**Authors:** Kaitlin A. Schaal, Marco La Fortezza, Lok Man Yuen, Gregory J. Velicer

## Abstract

Regulated aggregation and differentiation of cells is an intriguing form of multicellularity that produces diverse morphological forms, resulting from an interplay of complex evolutionary, regulatory, behavioral, and social mechanisms. We investigate the potential for predation to shape aggregative multicellularity evolution across different combinations of three species comprising a multi-trophic food web. The bacterium *Myxococcus xanthus*, which can develop into spore-filled fruiting bodies, is a mesopredator; the bacterivorous nematode *Pristionchus pacificus* is an apex predator, and the bacterium *Escherichia coli* is a shared basal prey for both predators. The number and morphology of *M. xanthus* fruiting bodies responds evolutionarily to nematodes, regardless of whether *E. coli* is present. *E. coli* alone with *M. xanthus* reduces both fruiting body formation and spore production, but the presence of nematodes eliminates those effects. *M. xanthus* lineages with an ancestral antibiotic resistance mutation evolved less overall, revealing strong historical contingency and suggesting potential tradeoffs between antibiotic resistance and responsiveness to biotic selection. Our results suggest that predation both of and by mesopredators has played important roles in the evolution of aggregative multicellularity and reveal complex inter-trophic evolutionary interactions in a relatively simple three-species food web.

## Introduction

The advent of clonal multicellularity (CM) is a Major Evolutionary Transition and is much studied (Stanley 1973; Bonner 1998; Ratcliff et al. 2012; Pentz et al. 2015; Herron 2016; Herron et al. 2022), but less attention has been given to understanding the origins, diversity, and evolutionary drivers of aggregative multicellularity (AM). Through AM, microbial cells that live independently for part of their life cycle come together to engage in and benefit from group behaviors. AM emerged multiple times, in both prokaryotes and eukaryotes, yet understanding of its evolutionary origins and what selective factors drive its diversification – potentially including being subject to predation – remains poor (Pentz et al. 2020; La Fortezza and Velicer 2021; Márquez-Zacarías et al. 2021; Herron et al. 2022; La Fortezza et al. 2022*b*, 2022*a*; Isaksson et al. 2025; Jahan et al. 2025). Here we examine and compare multiple biotic selective pressures hypothesized to be relevant to the evolution of AM, namely both predation of and predation by an aggregative microbe.

Many different types of organisms perform AM, suggesting that it has relevant adaptive benefits (La Fortezza et al. 2022*b*). The best-studied AM species are the amoeba *Dictyostelium discoideum* and the bacterium *Myxococcus xanthus,* the latter being the focus of this study. AM represents a way that organisms can overcome the biophysical limitations of single cells to increase their biological complexity. AM involves complex signaling repertoires that enable cells to encounter one another and regulate their coordinated behaviors, suggesting multi-step evolution and significant benefits to get there (La Fortezza et al. 2022*b*). The lack of a single celled bottleneck, an intrinsic feature of CM, makes AM susceptible to evolutionarily persistent cheating (Grosberg and Strathmann 2007); that it persists further suggests adaptive benefits. Most known forms of AM involve two components: cells assembling into a macro-structure, and differentiation of some cells into a reproductive/stress-resistant sub-population (*e.g.*, spores). This suggests that AM may have originated as a mechanism for responding to biotic threats and/or to survive unfavorable environmental conditions (La Fortezza et al. 2022*b*). The latter hypothesis is supported by the formation of stress-resistant spores within aggregate structures in many AM systems in the lab, whereas the former hypothesis has been less investigated.

Predation has long been studied as a driving force in the evolutionary transition to multicellularity (Stanley 1973; Bonner 1998; Boraas et al. 1998; Claessen et al. 2014; Pentz et al. 2015; Kapsetaki et al. 2016; Herron et al. 2019), and indeed predation has been shown to select for CM in the lab (Herron et al. 2019). Although it has been hypothesized that AM might also provide protection from predation for a variety of aggregative microbes (Kessin et al. 1996; Velicer and Vos 2009; Dahl et al. 2011; Pérez et al. 2011; DePas et al. 2014; Müller et al. 2015), direct study of how predators of AM microbes influence the evolution of their aggregative behaviors and morphologies has been lacking.

We address this gap with a tri-trophic-level evolution experiment involving: (i) the social soil bacterium *M. xanthus* (order Myxococcales, aka ‘myxobacteria’ (Shimkets et al. 2006*a*; Velicer and Vos 2009)), (ii) *Pristionchus pacificus*, a nematode capable of preying on *M. xanthus* (this study), (iii) and *Escherichia coli,* a bacterium that serves as prey to both *P. pacificus* and *M. xanthus* (Hong and Sommer 2006; Morgan et al. 2010). All are found in soils and may interact in their natural environment (Hong and Sommer 2006; Morgan et al. 2010). *M. xanthus* aggregates to form spore-filled fruiting bodies (FBs) upon starvation, as well as upon interacting with some other bacterial species (Kroos et al. 2025). *M. xanthus* is well known as a predator, swarming through soil in coordinated groups and preying on a wide range of other microbes, including gram-negative and gram-positive bacteria and fungi (Bull et al. 2002; Morgan et al. 2010; Mendes-Soares and Velicer 2013; Livingstone et al. 2017). *M. xanthus* presumably uses FB formation combined with sporulation for the purpose of better surviving low-nutrient conditions (Kroos 2017). However, it may also use FB formation to better survive predators in the environment. Selection imposed by such predators may have thus contributed to the origin and stunning diversification of fruiting-body morphologies observed among the myxobacteria (Shimkets et al. 2006*b*; Velicer and Vos 2009; La Fortezza et al. 2022*b*; Kroos et al. 2025), as well as among aggregative systems more broadly such as the dictyostelids (Bonner 2009).

Because *M. xanthus* and many other aggregative microbes (*e.g.*, *D. discoideum*) are mesopredators (*i.e.*, predators of other microbes as well as potential prey for higher-order predators), they are ideal for studying multi-step trophic webs in ecological and evolutionary experiments. *M. xanthus* predation on other bacteria has featured in multiple evolution experiments (MyxoEEs, see myxoee.org (Velicer et al. 2024)). In MyxoEE-4, for example, *M. xanthus* adapted its foraging ability in response to prey abundance (Hillesland et al. 2009). In MyxoEE-6, it both adaptively evolved as a predator and drove evolution of prey responses, including changes in virulence traits (Nair et al. 2019; Nair and Velicer 2021). In MyxoEE-3, prey presence and identity were found to indirectly shape the evolution of *M. xanthus* fruiting body phenotypes (La Fortezza et al. 2022*a*).

How mesopredators respond evolutionarily to higher-order predators and how such responses shape the evolution and ecology of food webs are longstanding questions that have nonetheless received relatively little direct experimental attention (Brodie and Brodie 1999; Abrams 2000; Urban 2013). Here we present an evolution experiment in which a bacterial predator – *M. xanthus* – functions as a mesopredator in a multi-step food web. This experiment was conducted to test effects of *M. xanthus* both i) carrying out and ii) being subject to predation on the evolution of *M. xanthus* features, including behavioral traits during interactions with both prey and apex predators. In this first evolution experiment with a bacterial mesopredator, we allowed only *M. xanthus* to evolve in order to simplify analysis of evolutionary causation. This initial study emerging from the evolution experiment focuses on phenotypic evolution of *M. xanthus* traits related to aggregative development.

Here we investigated whether the prey or the apex predator was the primary driver of AM phenotypic evolution, including the effect of each alone and in combination. Replicate populations of *M. xanthus* evolved in each of four selective regimes that differed only in the presence or absence of two non-evolving biotic partners – *E. coli* as the basal prey and *P. pristionchus* as the apex predator (Figure 1A). The ancestral *M. xanthus* strain underwent aggregative development after limited growth on the experimental medium, due to amino acid restriction. There was therefore the potential for the prey and/or apex predator to impose direct selection on features of *M. xanthus* development. CFcc limits *M. xanthus* growth through amino acid restriction (Bretscher and Kaiser 1978) without similarly limiting *E. coli* growth (Kreth et al. 2013) (Figure 1A). Importantly, after depleting growth substrate from the nutrient agar, *M. xanthus* underwent aggregative development within each weekly evolution cycle in the absence of any biotic partners, creating the potential for addition of the basal prey and/or apex predator to impose direct selection on developmental features. To address the effect of the prey and the apex predator on mesopredator evolution, we quantitatively compared several developmental traits – including fruiting body number, fruiting body density, and spore production – between evolved populations their ancestors, and among evolution treatments.

**Figure 1.**
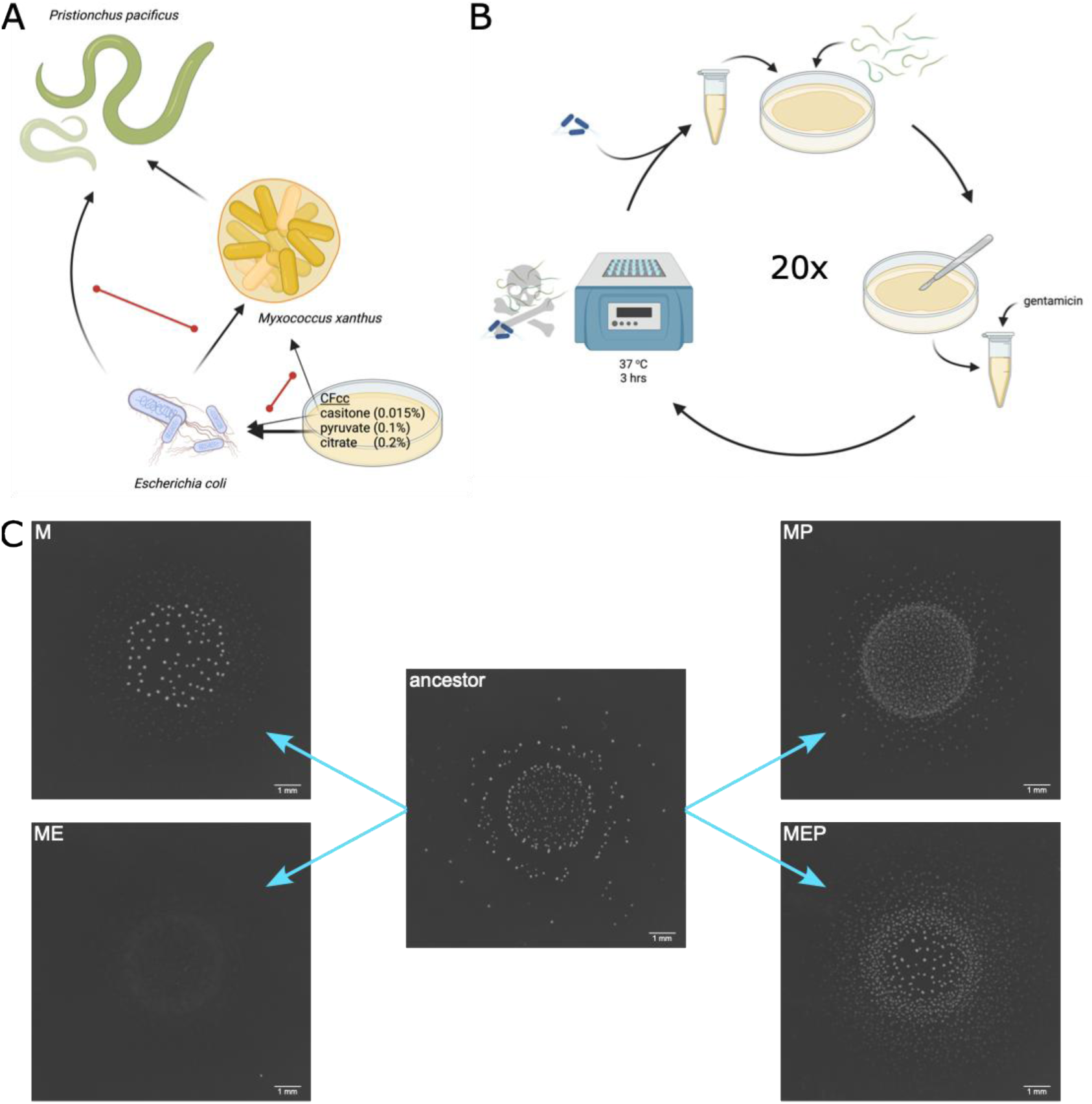
Experimental food web and evolution protocol. (A) We assembled a tri-trophic food web with *P. pacificus* as the apex predator, *M. xanthus* as the mesopredator, and *E. coli* as the basal prey.. Black arrows indicate energy flow (*e.g.,* via predation). Red double-headed lines indicate competition for resources. (B) Schematic showing the regime by which *M. xanthus* was transferred from cycle to cycle during the evolution experiment. (C) Images representing developmental diversification among evolved lineages. Images have been cropped, set to greyscale, and inverted for clarity. All images from the same ancestor. Panels A and B were created with BioRender.com.

## Methods

### Organisms

Our focal organism was *M. xanthus*, whose social traits and evolution in biotic context is of particular interest to us. Here we used the well-characterized *M. xanthus* lab strain GJV1 (Velicer et al. 2006) (strain ‘S’ in (Velicer et al. 1998)), as well as GJV2, a spontaneous mutant of GJV1 which is resistant to rifampicin (strain ‘R’ in (Velicer et al. 1998)). GJV2 differs from GJV1 by a single mutation in *rpoB*, the β-subunit of RNA polymerase (Zee et al. 2014). We used this antibiotic-resistant variant of GJV1 to initiate half of our populations to allow us to directly compete evolved populations against the reciprocally marked ancestor and distinguish them through selective plating during subsequent assays. We chose GJV2 because in previous studies it did not exhibit significant defects in either group swarming (Velicer et al. 2002) or development (Manhes and Velicer 2011), and we therefore did not expect substantial differences in trait evolution during trophic interactions.

Previous work indicates that *C. elegans* is not an effective predator of *M. xanthus* (Mayrhofer et al. 2021), so we used *Pristionchus pacificus* PS312 (Hong and Sommer 2006) as the apex predator. We observed that under laboratory conditions, *P. pacificus* is able to grow and reproduce when *M. xanthus* is given as the only food source (Figure 2). For the basal prey, we used *E. coli* OP50, which is a commonly used laboratory prey of nematodes (Brenner 1974).

**Figure 2.**
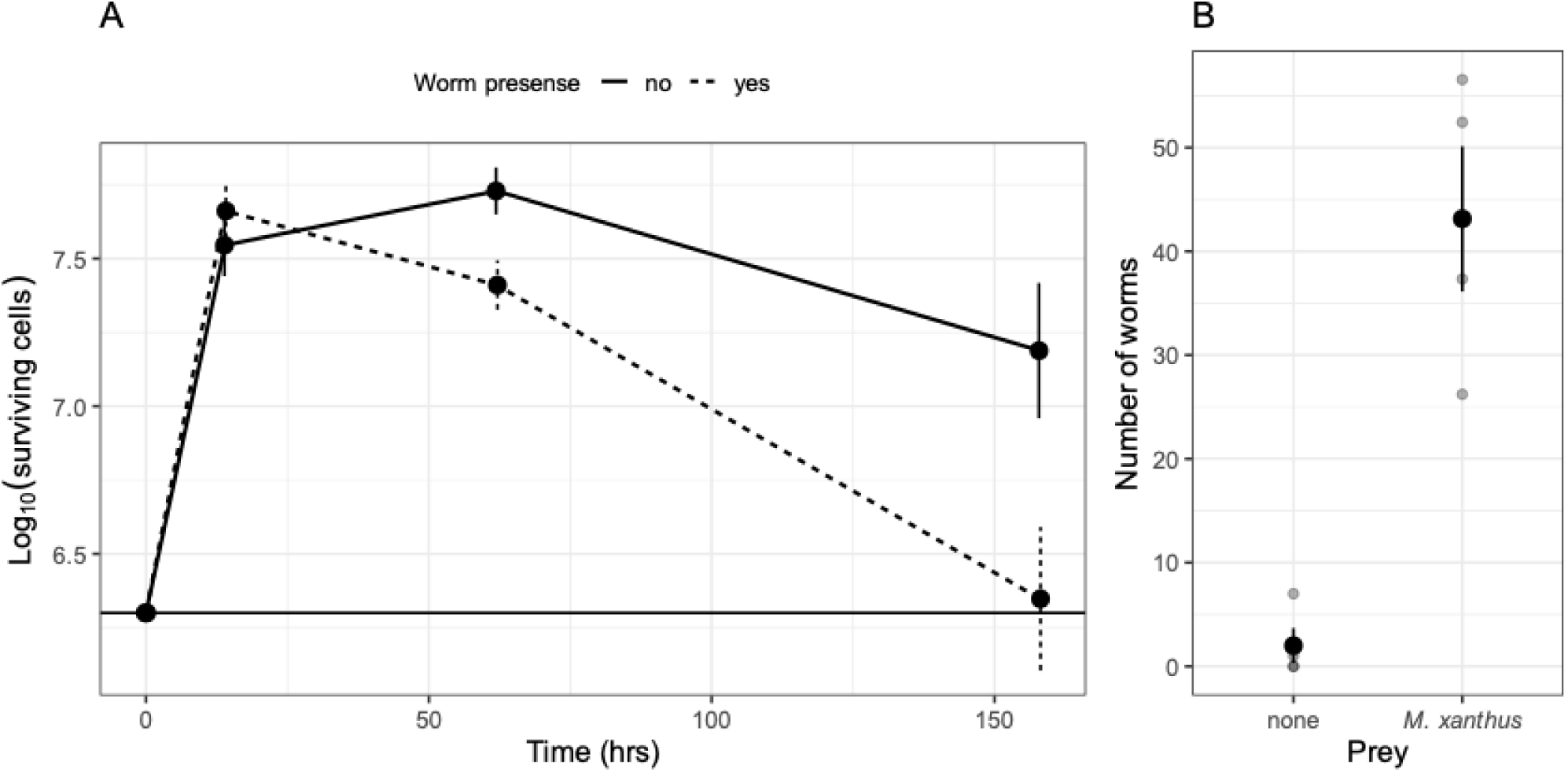
*Pristionchus pacificus* predation of *Myxococcus xanthus*. We cultured *P. pacificus* on *M. xanthus* as the sole food source. (A) Population dynamics of *M. xanthus* under predation by *P. pacificus*. Dots are averages of four biological replicates. The horizontal line shows the initial population size of *M. xanthus*. (B) *P. pacificus* reproduction on *M. xanthus*. Initial populations were founded with three J4 larvae. Large black dots are averages of four biological replicates. Error bars represent standard error. Small grey dots show individual replicates. *P. pacificus* population sizes after 72 hours are shown.

### Bacterial growth conditions

We maintained 20%-glycerol freezer stocks of bacterial strains at −80 °C. We inoculated *M. xanthus* strains onto CTT 1.5% agar plates (Bretscher and Kaiser 1978) and incubated at 32 °C and 90% rH for 4 days, then transferred a portion of the colony into CTT liquid and incubated overnight at 32 °C with shaking at 300 rpm until the cultures reached mid-log phase (OD_595_ 0.2 - 0.8). We inoculated *E. coli* strains directly into lysogeny broth (LB) and incubated overnight at 32 °C with shaking at 300 rpm.

### Culturing nematodes

We maintained freezer stocks of *P. pacificus* at −80 °C. We prepared freezer stocks according to Pires da Silva (Pires da Silva 2013). We collected worms from recently starved plates with Freezing Solution (100 µl/ml DMSO and 0.1 g/ml Dextran in water), aliquoted 1-ml volumes into freezer vials, and allowed them to freeze slowly in a Nalgene® Cryo 1°C Freezing Container containing isopropanol. To revive worms, we thawed an entire freezer vial, transferred the contents to a Falcon tube with a Pasteur pipette, added 10 ml of Thawing Solution (300 mg/L L-glutamine in M9 buffer [5 g/L NaCl, 6 g/L Na_2_HPO_4_, 3 g/L KH_2_PO_4_, and 1 mM MgSO_4_ in water]), centrifuged for 1 min at 150 x *g*, and removed the supernatant. We then transferred the thawed worms to the edges of *E. coli* OP50 lawns on 2% agar NGM (Stiernagle 2006) or HGM (Kauffman et al. 2011) plates. We incubated the plates at 25 C and 50% rH.

### Bleaching nematodes

We bleached *P. pacificus* as in Mayrhofer *et al*. (Mayrhofer et al. 2021) to create synchronized cultures to use in experiments. We washed nematodes from HGM plates using M9 buffer and collected them in a 15-ml Falcon tube. We centrifuged them for 1 min at 173 x *g* and removed all supernatant except 500 µl, to remove as much *E. coli* OP50 as possible. We added 1 ml of fresh M9 so that the total volume was 1.5 ml, then added 3 ml of bleaching solution (6 ml H_2_O, 6 ml NaOCl 5%, 2 ml 5M NaOH) and bleached with periodic vortexing until most of the adult bodies had dissolved, up to 7 minutes. We removed the bleach from the released eggs by centrifuging for 1 min at 173 x *g*, removing all but 500 µl of supernatant, and adding 4.5 ml sterile water. We washed the eggs four times, and after the fourth wash removed all but 500 µl of supernatant and added 3.5 ml M9. We used a Pasteur pipette to transfer the egg suspension to a 6-cm petri dish and added 1 µg/ml gentamicin to prevent any possible *E. coli* contamination. We allowed the eggs to hatch overnight. We removed dauer pheromone, which is released when food availability is low and inhibits nematode growth and development, and counted the number of hatched J2 larvae by centrifuging the worms in a 15-ml Falcon tube for 1 min at 173 x *g*, removing as much supernatant as possible, adding M9 up to 1 ml, and counting in triplicate 1-µl aliquots plated on sterile CFcc 1.5% agar plates (Mayrhofer et al. 2021).

### Nematode predation tests

To quantify population reduction of *M. xanthus* by *P. pacificus*, we prepared 6-cm petri dishes filled with 11 ml of CFcc 1.5% agar, inoculating ∼2 x 10^6^ CFUs GJV1 as the food source and spreading with glass beads. We added ∼200 J2 *P. pacificus* larvae in 100 µl M9 or 100 µl of sterile M9 as a predator-free control and incubated at 25 °C and 50% rH. After 1 day, 4 days, and 7 days we destructively harvested all biological material from agar surfaces with a sterile scalpel and transferred to the center of a 9-cm CTT 0.5% agar plate. We measured the diameter of the swarm size of the surviving *M. xanthus* population after 4 days and used a standard curve relating swarm size to inoculum size to estimate *M. xanthus* population sizes (approach adapted from (Rendueles and Velicer 2017), Fig. S5). We calculated the area under the curve and used a general linear model to test for differences with and without worms.

To quantify reproduction of *P. pacificus* using only *M. xanthus* as a food source, we inoculated 3 x 10^9^ cells of GJV1 or 300 µl sterile TPM buffer in the center of TPM 1.5% agar plates, picked 3 J4 *P. pacificus* larvae onto each plate, and incubated at 25 °C and 50% rH for 3 days. To count the worms, we washed the plates with 1 ml of TPM buffer, counted the number of worms present in 3 aliquots of 30 ul, and counted the number of worms remaining on each plate. For buffer plates, we only counted the worms present on the plates. We used a Mann-Whitney U-test to compare the two treatments.

### Evolution experiment

We isolated 6 clones each from GJV1 and GJV2 and named them 11-16 and 21-26, respectively (hereafter referred to collectively as ‘A’ for ‘ancestor’). We used each of these clones to found four populations, one in each of the evolution treatments. The evolution treatments were *M. xanthus* evolving in monoculture (M), *M. xanthus* evolving in co-culture with non-evolving *E. coli* (ME), *M. xanthus* evolving in co-culture with non-evolving *P. pacificus* (MP), and *M. xanthus* evolving in co-culture with non-evolving *E. coli* and non-evolving *P. pacificus* (MEP). We performed the experiment on CFcc 1.5% agar plates (Hagen et al. 1978; Mayrhofer et al. 2021), which contain 0.015% casitone as the only source of amino acids, as well as 0.1% pyruvate and 0.2% citrate. *M. xanthus* must exogenously acquire several amino acids (Bretscher and Kaiser 1978) and was therefore more limited in monoculture growth on experimental CFcc plates than *E. coli* OP50, a leaky uracil auxotroph (Brenner 1974). Both citrate and pyruvate stimulate *M. xanthus* growth in the presence of exogenously provided amino acids, with pyruvate stimulating greater growth (Bretscher and Kaiser 1978); once the casitone was depleted, *M. xanthus* would be unable to continue growing on the provided carbon sources.

To initiate the experiment, we added ∼2 x 10^6^ CFUs of each founding clone, pre-mixed with ∼10^9^ CFUs of *E. coli* for ME and MEP treatments, to 6-cm petri dishes filled with 11 ml of CFcc 1.5% agar. We spread the bacterial inoculum with glass beads until dry, removed the glass beads, and added ∼200 J2 *P. pacificus* larvae in 100 µl M9 (for MP and MEP treatments) or 100 µl of sterile M9 (for M and ME treatments). We sealed the plates with parafilm and incubated for 7 days at 25 °C and 50% rH.

To passage *M. xanthus* at the end of each cycle, we harvested the biological material from the surface of each evolution plate with a sterile scalpel and washed it into an Eppendorf tube with 1 ml of CFcc liquid with 1 µg/ml gentamicin to kill the *E. coli*. We incubated the tubes in a dry bath at 37 °C for 3 hrs (to kill the worms). We then transferred 1% of the volume of each culture to a new evolution plate (pre-mixed with either *E. coli* or buffer). We continued for a total of 20 cycles (Figure 1B). Six populations were lost due to contamination, leaving 42 evolved populations at the end of cycle 20.

At the end of even-numbered cycles, we transferred the remaining volume from the finished cycle to CTT liquid and allowed *M. xanthus* to grow up overnight in shaken culture at 32 °C and 300 rpm, then prepared 20%-glycerol freezer stocks. All following work was performed on population samples from terminal freezer stocks.

### Fruiting body morphology assay

We observed and quantified the fruiting body phenotypes of terminal evolved populations and their ancestral clones as in La Fortezza and Velicer (La Fortezza and Velicer 2021). We grew liquid cultures of cycle-20 evolved populations as described above. We diluted them and allowed them to shake overnight again in order to ensure that the resulting cultures would be as free from clumps as possible. We centrifuged them at 12,000 rpm for 5 min and resuspended to 5 x 10^9^ CFUs/ml in TPM buffer. We poured 5 ml of CFcc 1.5% agar into 6-cm petri dishes and allowed to dry. We plated 10 µl of each population and allowed the spots to dry for 1 hr, then incubated at 25 °C and 50% rH for 7 days. Due to limitations on sample handling time during assay plate preparation, we processed in 5 blocks per replicate of 11 populations at a time. Each block also included GJV1 and GJV2 as internal controls. This resulted in 13 samples per block, and 20 total blocks.

After 7 days, we imaged the samples (Figure 1C) using an Olympus SZX16 microscope and Olympus DP80 camera together with cellSens software version 1.15 (Olympus, Tokyo, Japan), according to La Fortezza and Velicer (La Fortezza and Velicer 2021). Microscope, camera, and software settings were the same for all samples across all blocks and replicates. Images are available via the BioImage Archive (DOI). To check for correlations between fruiting body morphology and spore production, we then harvested all biological material using flame-sterilized scalpels, sonicated to disperse aggregates, and quantified spore production by counting the number of spores (identified by spherical shape and high refractivity) in a given area of a Neubauer improved hemocytometer.

### Image processing

We analyzed the images using Fiji version 1.53q (Schindelin et al. 2012). We converted each image to 8-bit, removed dust particles and other non-bacterial parts of the image, and set the black-white threshold to capture all aggregates. We saved this information as an overlay which we applied to the original image to collect data on the aggregates. We quantified fruiting body morphology with four traits previously used for this purpose (Rivera-Yoshida et al. 2019; La Fortezza and Velicer 2021): number of aggregates, size of each aggregate (pixels^2^), average grey value of each aggregate (density in La Fortezza and Velicer 2021), standard deviation of grey values within each aggregate (density heterogeneity in La Fortezza and Velicer 2021), and x and y coordinates of each aggregate. The 1 mm scale bar (Figure 1C) corresponds to 293 pixels; 1 pixel = ∼3.5 nm. Fruiting body density is inversely related to transparency and positively correlated with fruiting body maturity and spore content; here we use an arbitrary scale between 0 and 200. Density heterogeneity quantifies within-aggregate texture variation. A full description of these traits is given in (Rivera-Yoshida et al. 2019; La Fortezza and Velicer 2021).

### Fruiting body trait analysis

We analyzed fruiting body morphological traits and spore production in R version 4.1.3 (R Core Team 2024) and RStudio version 2022.02.1+461 (RStudio Team 2020). We analyzed overall fruiting body morphology according to the method established by La Fortezza *et al*. (La Fortezza and Velicer 2021), by principal components analysis followed by PERMANOVA using the *adonis2* function in the “vegan” package (Oksanen et al. 2020) to check for effects of evolution environment. All trait data (fruiting body number, area, mean grey value, and density heterogeneity) were standardized (centered and scaled to unit variance) prior to PCA using the R command *prcomp()*, and we tested for homogeneity of multivariate dispersions using the *vegan::betadisper()* function. We further analyzed fruiting body traits individually using ANOVA to test for effects of evolution environment and of ancestral genotype (GJV1 or GJV2, from which we picked the 12 founding clones used to start the evolution experiment) followed by Tukey HSD tests for differences from the founding clones and among evolution treatments. We analyzed spore production data the same way, ANOVA for differences based on evolution environment and GJV1 versus GJV2 ancestry followed by Tukey HSD tests. Data occasionally violate normality or homoscedasticity. Non-parametric models (Kruskal-Wallis tests with Conover-Iman post-hoc tests) produce similar results to parametric ones and are less conservative; here we report results from parametric models to allow comparison of treatment means. The data files and full statistical analysis are available via Zenodo (DOI). There were different numbers of surviving populations per evolution treatment and *rpoB* ancestry. For GJV1 lineages, M = 5, ME = 6, MP = 5, MEP = 4. For GJV2 lineages, M = 5, ME = 6, MP = 5, MEP = 6. We replicated the post-evolution development assay 4 times, resulting in 4 replicates per lineage, and therefore ≥16 assay values per evolution treatment. Figures were generated using the “ggplot2” package (Wickham 2016).

### Morphological integration analysis

We tested morphological integration of fruiting body traits by analyzing eigenvector variance of the trait morphospace of the four developmental traits following previously established methods (Pavlicev et al. 2009; Machado et al. 2019; La Fortezza et al. 2022*a*). Traits were standardized prior to computing covariance matrices. For this analysis, we included spore production as a trait. We calculated covariance matrices of the five traits for each evolution treatment individually and eigenvector variances (Var(λ)) using *CalcEigenVar()* in the “evolqg” package (Melo et al. 2016). Figures were generated using the “ggplot2” package (Wickham 2016).

### Time-lapse assay

We performed a time-lapse fruiting body morphology assay on GJV1-descended lineages from cycle 20 developing on CFcc media, initiated as described above. We imaged the samples using the Olympus SZX16 microscope and Olympus DP80 camera with cellSense software version 4.3 (Olympus, Tokyo, Japan) every 24 hours for seven days. The settings were kept identical throughout all replicates (exposure time = 1.67ms, lens = Olympus 0.5x PF, zoom = 1.25, ISO = 200, illumination = BF built-in system). We performed four independent replicate observations of each lineage.

We processed and analyzed the images with Fiji version 1.54p, using the same technique as the described above with the addition of excluding all particles under 30 pixel^2^ before overlaying onto the original image. For samples without fruiting bodies, we set the number of aggregates to one, average grey value to 50, the mean standard grey value to 0.18, and the area of the aggregate to 30 pixel^2^. This corresponded to one very small, pale and homogeneous fruiting body on the entire plate. We generated an artificial starting point (t = 0 hour) using the same values to improve readability of the graphs. We performed a principal component analysis as above using R version 4.1.3 (R Core Team 2024) and RStudio version 2022.02.1+461 (RStudio Team 2020), and generated figures using the “ggplot2” package (Wickham 2016).

## Results

*P. pacificus successfully preys on M. xanthus.* To confirm whether *P. pacificus* acts as an apex predator in this system, we tested its ability to reproduce and consume *M. xanthus* as a sole food source. *P. pacificus* significantly reduces *M. xanthus* population size (Figure 2A; one-way ANOVA for area under the curve, *F_1,20_* = 6.94, *p* = 0.039, difference = 86.5% reduction at final time point), and is able to complete development to adulthood and reproduce successfully (Figure 2B; Mann-Whitney U-test, n = 4, W = 0, *p* = 0.02, difference = 41 worms).

*rpoB-mutant lineages show far less trait evolution in communities than wild-type lineages.* We anticipated change in phenotypes related to multicellular development over evolutionary time in all treatments, as well as phenotypic diversification as a function of treatments, as such traits are relevant to nutrient levels, prey presence and type (La Fortezza et al. 2022*a*), and we hypothesized also to anti-predator defense (La Fortezza et al. 2022*b*). We therefore examined four traits related to fruiting body morphology in the evolved populations and their ancestral clones: the number of fruiting bodies produced by each population, their average size, their average density (darkness of color, with darker aggregates generally being more mature and containing more spores), and the average heterogeneity of density.

In the evolution experiment we included six lineages per treatment descending from GJV1 (wild-type) and six descending from GJV2 (spontaneous rifampicin-resistant *rpoB* mutant), to allow the possibility of competing oppositely-marked strains head-to-head during follow-up assays. We chose this design because these strains have been used this way before (Velicer et al. 2002; Manhes and Velicer 2011). However, observations during the evolution experiment indicated that lineages founded from the two strains showed different patterns of evolution. We therefore performed an initial analysis of the principal components of variation among the four morphology traits based on ancestral genotype (GJV1 or GJV2), including the 12 ancestral clones and all surviving populations from the four evolution treatments from the final timepoint (N = 26 for GJV1, N = 28 for GJV2). In the resulting PCA plot (Figure 3A), descendants of GJV2 cluster more closely together, indicating less evolutionary divergence from the ancestral clones and among evolution treatments than for lineages descending from GJV1. Although previous studies found no significant differences compared to wild-type swarming and development phenotypes (Velicer et al. 2002; Manhes and Velicer 2011), the *rpoB* mutation likely affected GJV2 descendants’ ability to evolve in response to evolutionary conditions. We therefore analyzed principal components of trait variation for GJV1 and GJV2 descendants separately.

**Figure 3.**
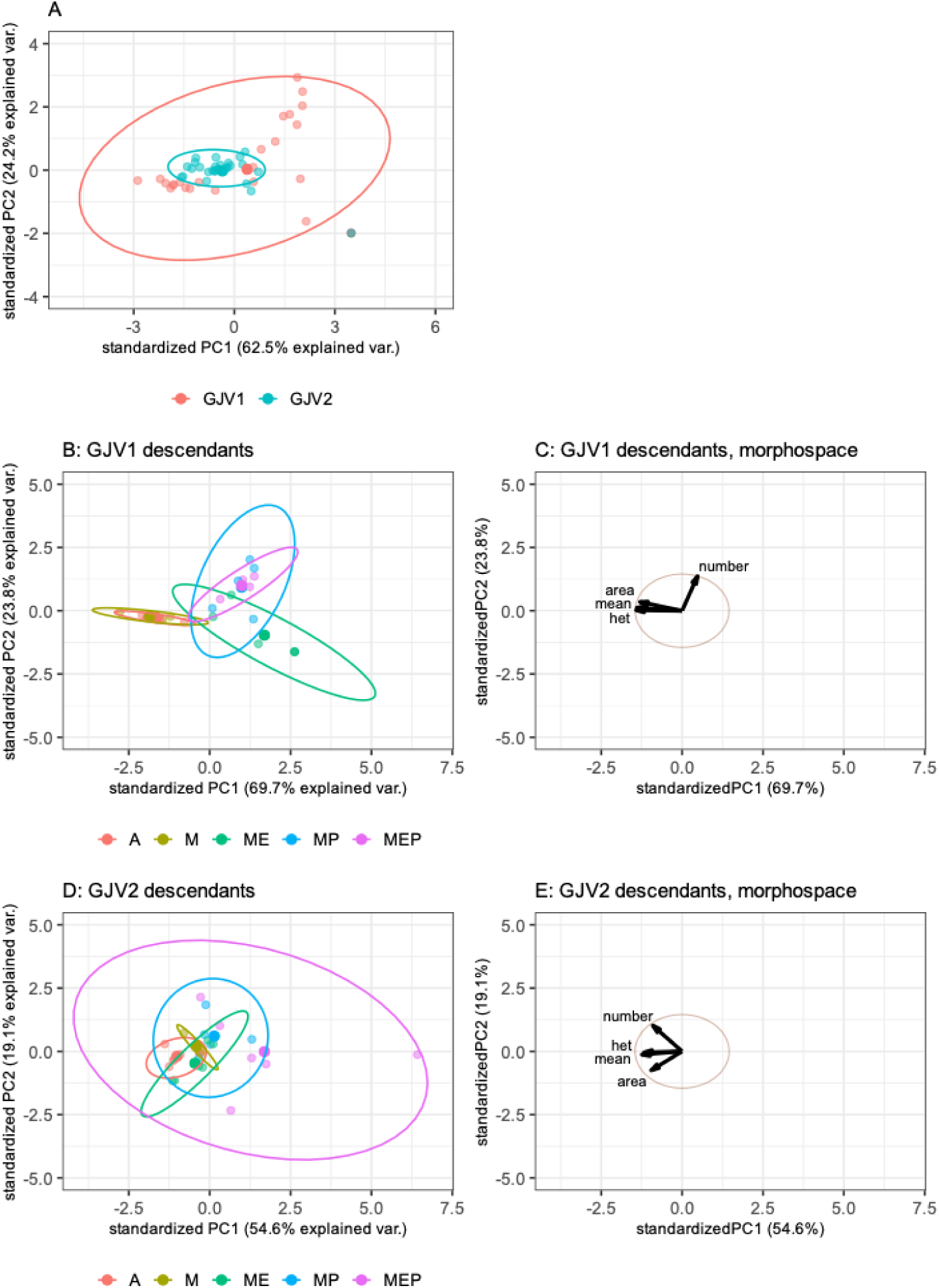
Evolutionary variation depends on ancestral genotype. We analyzed morphologies of fruiting bodies produced by *M. xanthus* populations that had evolved alone (M), in the presence of *E. coli* (ME), in the presence of *P. pacificus* (MP), or in the presence of both (MEP), as well as the clones used to found the evolved populations (A). Six of the founding clones come from GJV1 and six come from GJV2. We measured the principal components of variation across fruiting body traits. Here we show overall variation in fruiting body morphology as a function of the ultimate ancestor of the clones or evolved populations in the form of a principal components analysis of the four traits. Panel A shows the phenotypic variation within descendants of GJV1 versus descendants of GJV2. Panels B & D show trait variation based on evolution environment. Small dots represent averages across four replicates for each surviving lineage, large dots represent the centroid of each group (the average of PC1 and PC2 across all replicate lineages) and ellipses represent the 95% confidence region. Percentages listed in the x- and y-axis titles are the percent of variation within the dataset which is explained by that principal component. Panels C & E visualize the morphospaces, which consider how the four traits correlate, and which can be used to interpret the variation seen in panels B & D. Arrows indicate the directional effect of each trait on the morphospace – along that vector, the value of the indicated trait increases. Traits: “het” = fruiting body heterogeneity, “area” = fruiting body size, “mean” = fruiting body darkness, “number” = number of fruiting bodies produced by a defined population of cells.

GJV1 lineages show significant variation among the five groups (evolved populations and ancestral clones; Figure 3B; PERMANOVA for effect of treatment, *F*_4,21_ = 16.99, *p* < 0.001). GJV2 lineages also vary (Figure 3D; PERMANOVA for effect of treatment, *F*_4,23_ = 3.25, *p* = 0.002), although trait differences are smaller in magnitude than for wild-type lineages (Figure 4). Further, we see different patterns of evolved integration, or pleiotropic correlation, across fruiting body morphology traits (Figure S1).

**Figure 4.**
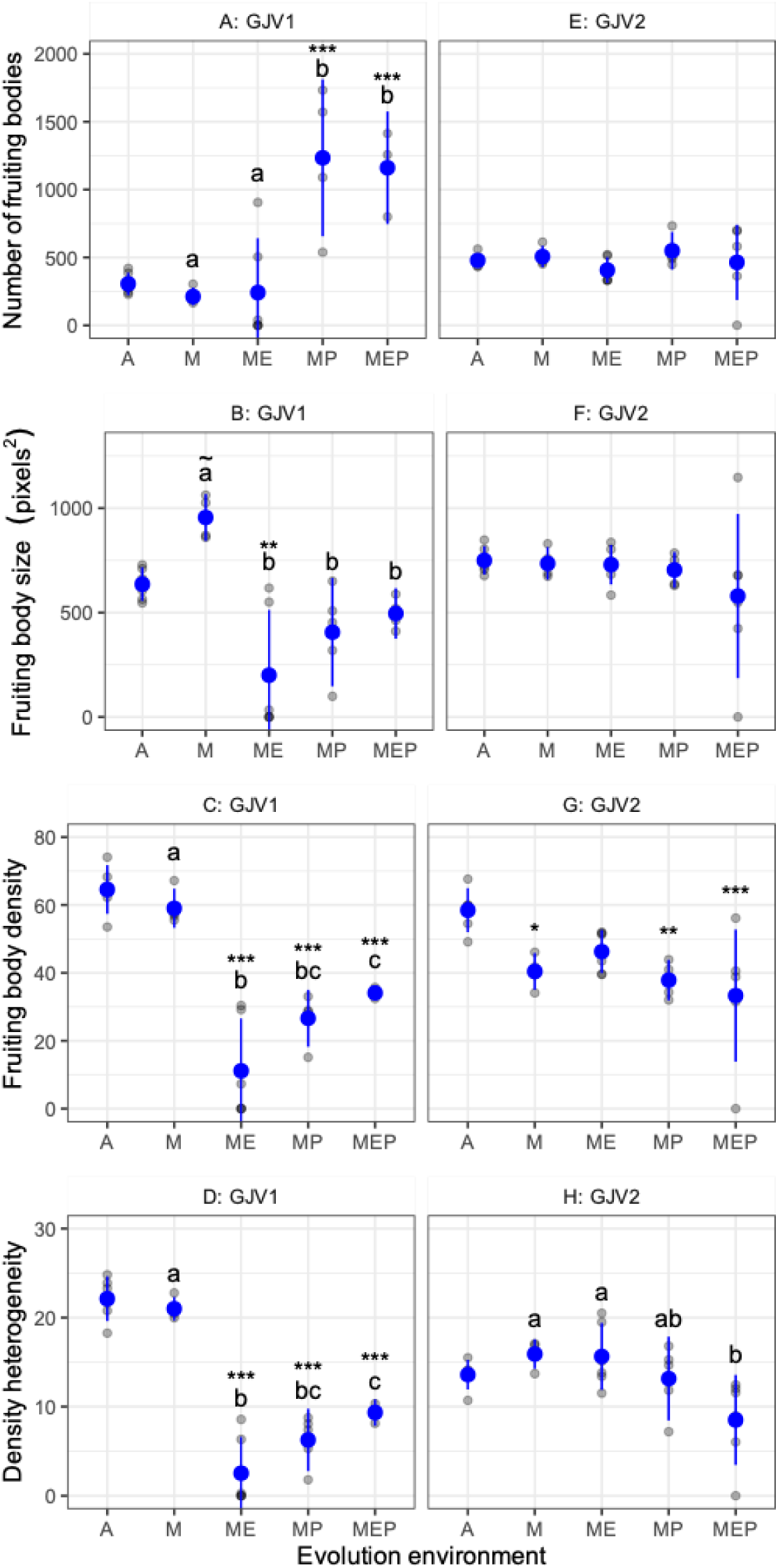
Fruiting body traits show evolutionary responses to community structure. Each measured trait of fruiting body morphology – number, size, density, and density heterogeneity – is plotted individually. We observe differences from the ancestral phenotype, represented by the founding clones (A on the x-axes) and differences among evolution treatments. Small grey dots show the mean of each evolution lineage across 3-4 biological replicates, large blue dots represent the mean across 4-6 replicate lineages, and blue bars show 95% confidence intervals of cross-lineage means. Asterisks indicate difference from the ancestors: ∼ = *p* < 0.06, * = *p* < 0.05, ** = *p* < 0.01, *** = *p* < 0.005. Letters indicate statistical groups of evolution treatments that do not differ within groups but differ between groups (*p* < 0.05). Vertical panels group clones or populations descending from GJV1 or from GJV2.

*The apex predator drives evolution of fruiting body number in GJV1 descendants*. Two striking results emerged from the PCA of the fruiting body morphologies of GJV1 descendants. First, we see clear divergence of all three treatments with multi-species communities (ME, MP and MEP); Figure 3B, green, blue and purple ellipses, respectively) from the founding clones (A) and the M populations (Figure 3B red and yellow ellipses, respectively). Second, we also observe clear divergence of the two multi-species treatments that included nematodes (MP and MEP) from the one that did not (ME).

Quantifying individual traits (Figure 4), we see that MP and MEP populations produce more fruiting bodies that are also smaller, less dense, and less heterogenous, while the founding clones and the M populations produce fewer fruiting bodies that are relatively larger, denser, and more heterogenous (Figure 1C, representative images). In greater detail, populations which evolved in the presence of *P. pacificus* (MP and MEP) show significantly higher numbers of fruiting bodies than the founding clones or the M or ME lineages (Figure 4; Tukey HSD tests, *p*’s < 0.002, differences > 855). This effect is independent of the presence of *E. coli* (Tukey HSD test, *p_MP:MEP_* = 0.99, difference = 73). Fruiting body number of M and ME treatments did not diverge from the ancestral phenotype (Figure 4; Tukey HSD tests, *p*’s > 0.98). GJV2 descendants do not show a similar trend of phenotypic change (Figure 4; one-way ANOVA for effect of treatment, *F*_4,23_ = 0.76, *p* = 0.56). Together, this indicates that increasing the number of fruiting bodies produced by a population of a given size is an evolutionary response specific to the presence of worms, but that the *rpoB* mutation in the GJV2 lineages inhibits this response.

We hypothesized that MP and MEP lineages produced more, smaller fruiting bodies because they entered development earlier than other lineages, possibly as a defensive adaptation to the presence of *P. pacificus*. To test this hypothesis, we conducted a time lapse experiment, allowing all lineages to develop as in the previous assay and imaging them at 24-hour intervals.

While MP and MEP lineages show differing developmental trajectories compared to the other treatments, as expected from the above results, they do not show evidence of earlier initiation of development (i.e., increased distance of day 1 from t=0 than other treatments; Figure 5A). The number of fruiting bodies observed in MP and MEP lineages generally increases between days 1 and 3 after initiation of the experiment, whereas the fruiting bodies of the ancestral clones (A) seem to have developed within the first 24 hours (Figure 5B). Overall, these results indicate that the increased number and smaller size of MP and MEP fruiting bodies is not a result of the population initiating development earlier, before cells have had a chance to aggregate in greater numbers.

**Figure 5.**
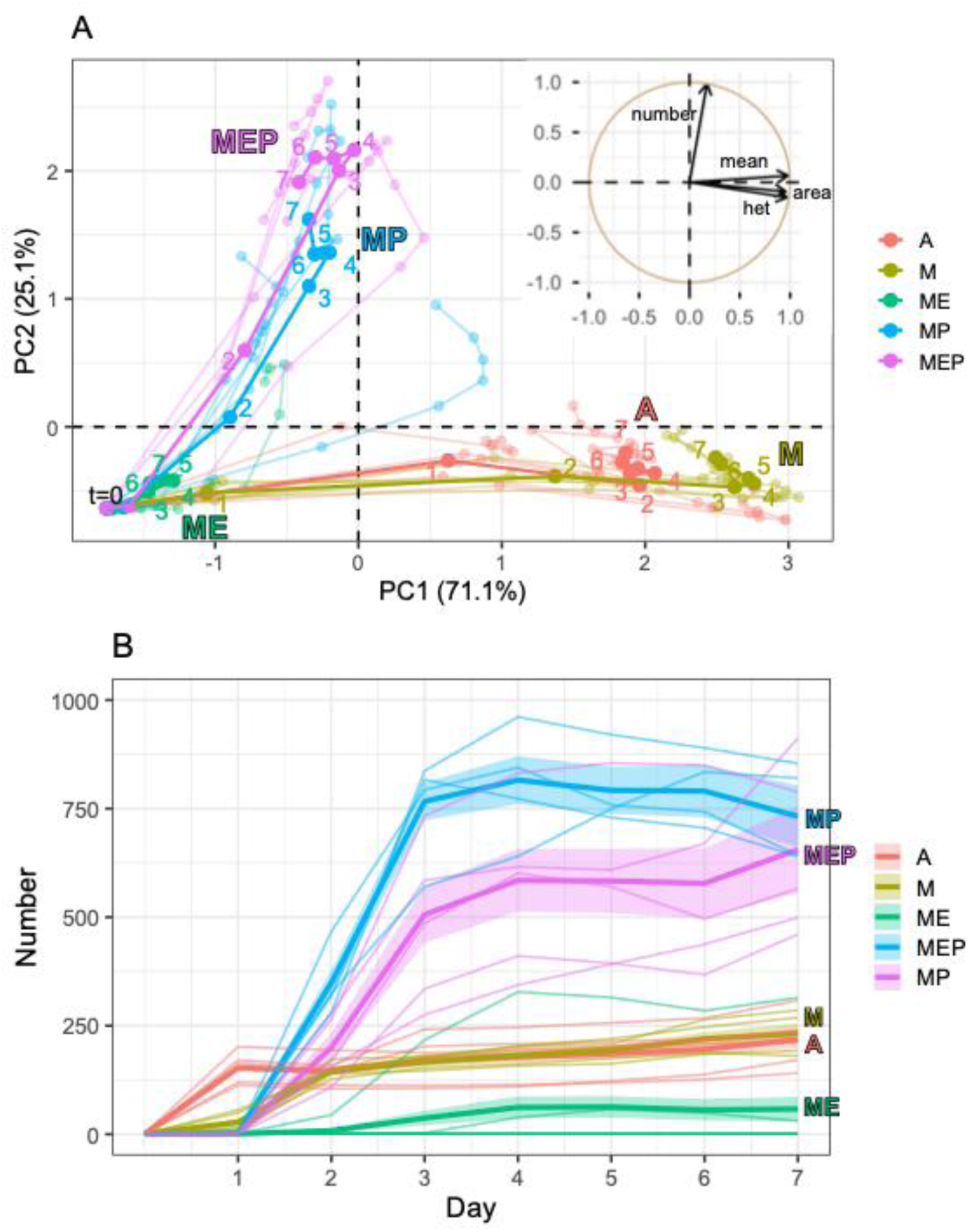
Development over seven days. We imaged development of evolved *M. xanthus* populations over seven days. (A) PCA trajectories of fruiting body morphology. Solid dots and lines show averages per treatment. Transparent dots and lines show individual lineages, averaged over four biological replicates.

*Evolution in multi-species communities reduces individual-fruiting-body trait values and promotes morphological trait integration*. The PCAs in Figure 3B and 3D suggest little overall divergence between phenotypes of populations that evolved in monoculture and phenotypes of the ancestral clones (*i.e.,* the red and yellow ellipses are largely overlapping, indicating that they represent groups with similar trait values). In line with this, the individual trait analyses (Figure 4) show that the M populations evolved to differ from the ancestral phenotype only in fruiting body size, and not at any of the other three traits. The average fruiting body size of the M populations increased significantly relative to the average values for the other evolution treatments (Tukey HSD tests, *p_M:ME_* < 0.0001 difference = 755 pixels^2^ (2.58 mm^2^), *p_M:MP_* = 0.0009 difference = 550 pixels^2^ (1.88 mm^2^), *p_M:MEP_* = 0.009 difference = 460 pixels^2^ (1.57 mm^2^)), although the difference relative to the ancestor was not significant (*p* = 0.06, difference = 319 pixels^2^ (1.09 mm^2^)). The average size of ME fruiting bodies decreased relative to the ancestor (Tukey HSD test, *p* < 0.004 difference = 436 pixels^2^ (1.49 mm^2^)), although this is most likely due these populations no longer producing mature fruiting bodies.

Contrary to the increase in fruiting body number seen in MP and MEP populations descending from GJV1, there are broad decreases in other trait values across all three treatments with multi-species communities (Figure 4). GJV1 descendants that evolved under ME, MP, or MEP conditions all form significantly less dense and less heterogeneous fruiting bodies than both the ancestor (Tukey HSD tests, *p*’s < 0.0002, grey value differences >30, heterogeneity differences >12) and the M-evolved populations (Tukey HSD tests, *p*’s < 0.003, grey value differences >24, heterogeneity differences >11). In addition, ME fruiting bodies are less dense than MEP (Tukey HSD test, *p* = 0.004, difference = 23). Both metrics refer to pigmentation of the fruiting bodies, measured as grey value of the pixels arbitrarily scaled from 0 to 200. Higher density correlates with more mature fruiting bodies containing greater numbers of spores; higher density heterogeneity indicates greater texture variation on the outer surface of the fruiting body. That these traits did not also decrease among the M populations points to evolutionary responses specific to the presence of biotic partners rather than a generic response to abiotic conditions.

Evolution in ME, MP, and MEP conditions increased covariance of fruiting body traits. To investigate general trends in morphological trait integration related to evolution environment, we calculated eigenvector variances (Var(λ)) for the four traits plus spore production for the four environments and the ancestral clones. Var(λ) of 0 indicates no observed correlation among traits, while Var(λ) of 1 indicates that most variance among traits is explained by the first principal component, suggesting stronger covariation (Pavlicev et al. 2009). The ancestral state has eigenvector variance of 0.5, showing partial correlation among the morphological traits. This increases for M, and increases to nearly 1 for ME, MP, and MEP (Figure S1). This suggests that evolution in multi-species communities caused *M. xanthus* developmental traits to evolve in a more integrated manner than did evolution in single-species populations.

Populations descending from GJV2 also showed some evolution at traits other than fruiting body number, but changes were smaller overall than for GJV1-descendants (as expected from the PCA in Figure 3A). All evolution treatments except for M form less-dense fruiting bodies than the ancestor (Tukey HSD tests, *p*’s < 0.05, differences >18). MEP populations show divergence in density heterogeneity from M- and ME-evolved populations (Tukey HSD tests for differences among evolution environments, *p*’s < 0.02, differences >7).

*Interaction between the apex predator and basal prey causes unexpected phenotypic evolution*. If the selective effects of the basal prey and the apex predator on *M. xanthus* phenotypic evolution were additive, we would expect trait values in the MEP treatment, which exposed *M. xanthus* to both, to fall in between those for ME and MP treatments. However, instead we see striking deviations from this expectation for two individual fruiting body traits – density and density heterogeneity (Fig. 4C-D). If one starts with MP as a reference point and then considers that the ME result would predict that the MEP value should be lower than the MP value under the assumption of linearity, we see that inclusion of *E. coli* in the MEP populations has the opposite effect, causing these three trait values to evolve to higher levels than in the MP populations rather than lower. And using ME as the starting reference point, inclusion of *P. pacificus* in the MEP treatment increases the trait values more than predicted from the ME and MP data alone. Thus, the effects of *E. coli* and *P. pacificus* on *M. xanthus* evolution interact in a nonlinear manner.

Numbers indicate the days since the start of the experiment. The trajectory starting point (t=0) was calculated as described in the Methods. Inset shows trait morphospace. Traits: “het” = fruiting body heterogeneity, “area” = fruiting body size, “mean” = fruiting body darkness, “number” = number of fruiting bodies produced by a defined population of cells. (B) The number of fruiting bodies produced over time. Thicker lines show treatment averages, with ribbons indicating standard error. Thinner lines show individual lineages, averaged over four replicate observations.

*Nematodes prevent evolutionary reduction of M. xanthus sporulation caused by E. coli alone*. Given the significant changes in fruiting body numbers and several traits of individual fruiting bodies, such as size, we hypothesized that there may be correlated changes in the number of spores produced by those populations and associated treatment-level effects. We paired each measure of fruiting body number and size with the number of spores counted from that plate (Figure S2). Spore production correlates positively with both of these traits (Table S1). For descendants of GJV1, the correlation with fruiting body size differs based on evolution environment (Table S1; two-way ANOVA for effects of area and treatment on spore production, *F*_4,16_ = 3.132, *p* = 0.044), but in every other case the effects are independent.

When looking at spore production only (Figure 6) overall across populations descended from both ancestors (GJV1 vs GJV2), most treatments showed no significant change or moderate increases in spore production. Exceptionally, however the ME populations decreased ∼90% in spore production on average. The absence of a corresponding decrease in the MEP lines indicates that the presence of *P. pacificus* prevented selective effects of *E. coli* from evolutionarily degrading *M. xanthus* sporulation, possibly due to the removal of *E. coli* by nematode predation.

**Figure 6.**
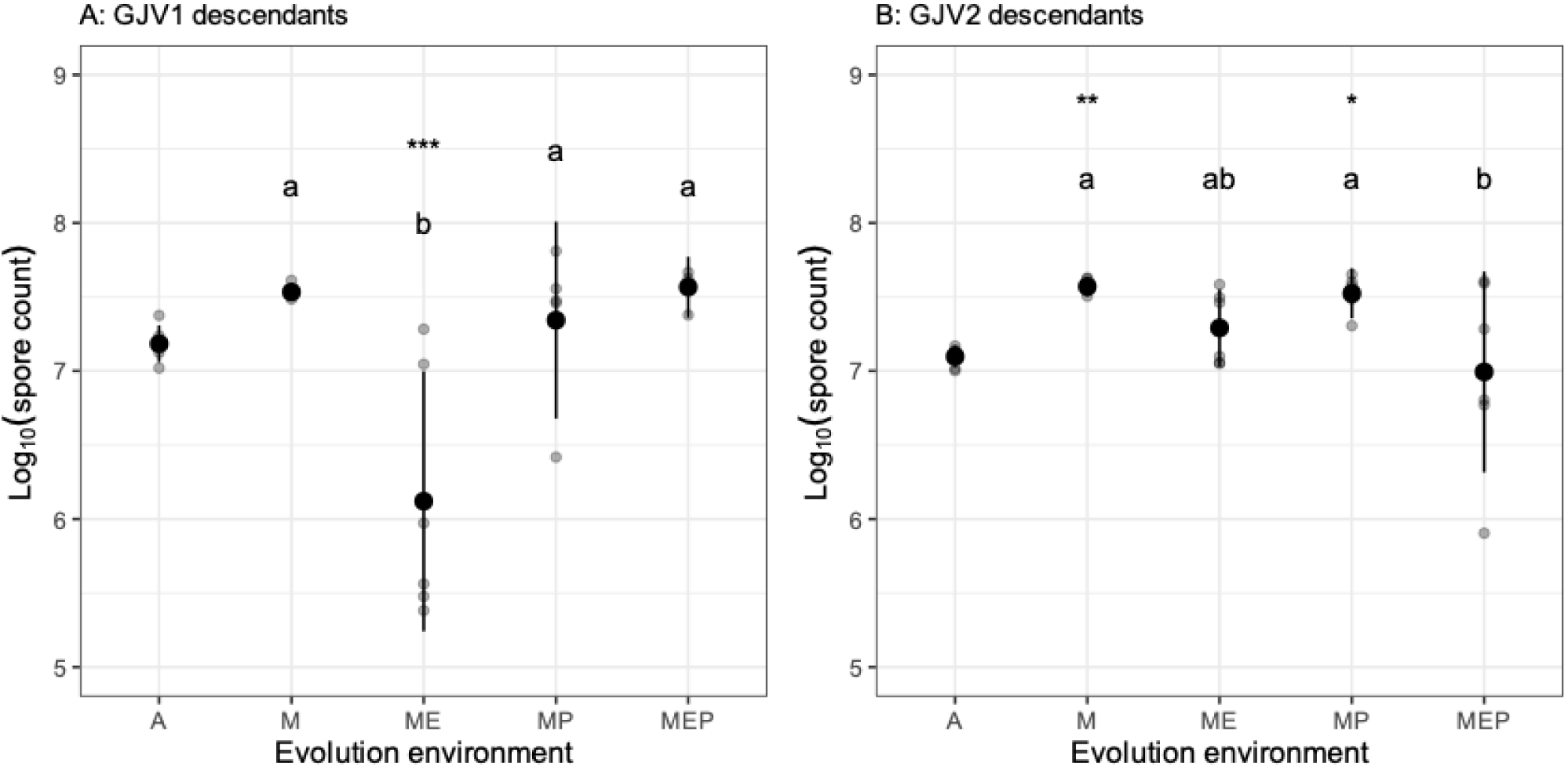
Spore production by GJV1 and GJV2 descendants. We counted the number of spores produced on each fruiting body morphology analysis plate. Small grey dots show the mean of each evolution lineage across 3-4 biological replicates, large black dots represent the mean across 4-6 replicate evolutionary lineages, and blue bars show 95% confidence intervals of cross-lineage means. Asterisks indicate difference from the ancestors: * = *p* < 0.05, ** = *p* < 0.01, *** = *p* < 0.005. Letters indicate statistical groups of evolution treatments that do not differ within groups but differ between groups (*p* < 0.05).

## Discussion

Here we characterize the phenotypic evolution of an aggregatively multicellular bacterium in a multi-trophic food web, with implications for understanding the evolution of biotic networks and aggregative developmental systems, including roles of historical contingency. The presence of a nematode predator of *M. xanthus* strongly impacts the evolution of both *M. xanthus* fruiting body morphology and spore production. *P. pacificus* had several evolutionary effects on *M. xanthus*, including (i) increased fruiting-body numbers after evolution in both the absence and presence of *E. coli* (Figure 4A; MP-vs-M and MEP-vs-ME), (ii) reduced fruiting body size, density, and density heterogeneity during evolution in the absence of *E. coli* (Figure 4B-D; MP-vs-M), (iii) mitigation of larger decreases in fruiting body density and density heterogeneity caused by evolution with *E. coli* alone (Figure 2C,D; MEP-vs-ME, and (iv) prevention of major decreases in spore production caused by evolution with *E. coli* alone (Figure 5A; MEP-vs-ME). Increased trait integration (Figure S1) suggests selection on development (Pavlicev and Hansen 2011).

### Apex predation shapes aggregative developmental evolution

We show that the basal prey and the apex predator in our experimental system in some cases had an interacting influence on trait changes in the mesopredator. Evolution in the presence of prey and absence of predators often led to the lowest values of morphological traits (Figure 4) and spore production (Figure 5) (La Fortezza et al. 2022*a*). In contrast, evolution in the presence of *P. pacificus* tended to lead to greater numbers of fruiting bodies relative to the ancestral phenotype (Figure 4A) and contributed to a trend of increasing density and density heterogeneity compared to the treatment with *E. coli* (Figure 4C-D). In the latter case, the presence of both organisms led to evolution of higher trait values than the presence of *P. pacificus* alone. This could indicate an effect like apparent competition between the two bacteria, where the basal prey boosts the apex predator’s population, increasing the apex predator’s effect on the mesopredator (Holt 1977; Holt et al. 1994; Holt and Bonsall 2017), or like trait-mediated indirect interactions, where pairwise species interactions are altered when one or both species respond phenotypically to the presence of a third (Werner and Peacor 2003). Collectively, the patterns of trait change show clear evidence of evolution in response to an apex predator which affects developmental pathways in such a manner that phenotypic changes emerge across different environments, in both 2- and 3-level trophic communities. Here we found no evidence for faster development in MP and MEP lineages, but further research into potential regulatory changes can investigate whether there are other treatment-level effects on developmental timing. Studies of the ecological parameters of the tripartite interaction considered here may provide insight into additional ecological drivers relevant here, such as the roles of effective population size *N_e_* in evolutionary trajectories across treatments and of non-consumptive effects of predators on prey populations, behavior, and phenotypes (Sheriff et al. 2020). Assessment of relative biomass of the three organisms may allow better understanding of how the ecological dynamics shape the food web functioning as an ecosystem (as in (Elmhagen et al. 2010)).

### Historical contingency constrains evolutionary responses to biotic selection

Observations during the evolution experiment led us to test how a single mutational difference between ancestral founding populations profoundly impacted *M. xanthus* evolvability in response to both predator and prey. Populations which descend from GJV2, an ancestor carrying a single-base-pair mutation in *rpoB*, underwent far less morphological change and diversification overall than populations lacking the mutation (Figures 3, 4, & S2). A previous study found differences between evolved populations descending from GJV1 and those descending from GJV2 in terms of colony-color evolution (Rendueles and Velicer 2017), showing additional evolutionary effects of GJV2’s *rpoB* allele. Single mutations have been observed to determine fitness trajectories in bacteria (Aggeli et al. 2021). Our findings contribute to the growing understanding of how very few genetic differences can profoundly influence how lineages respond evolutionarily to the same selective conditions in unexpected ways, and relatedly, how ultimate evolutionary outcomes can be contingent on small initial genetic differences (Blount et al. 2008, 2018). They also point to great complexity in interplay of forces shaping the evolution of AM mesopredator systems such as *D. dictyostelium* and *M. xanthus,* including not only who they encounter as their predators and prey, but also evolutionary consequences of evolving resistance to antagonistic biotic compounds such as antibiotics. The profound evolutionary consequences of this single mutation in our study may be due to the mutation causing systematic differences in what adaptive pathways are followed or to differences in the effects of similar adaptive pathways on the observed developmental phenotypes due to epistasis (Blount et al. 2018; Aggeli et al. 2021). Alternately, the different evolutionary outcomes between GJV1 and GJV2 may be due to differences in effective population size *N_e_*, which is expected to constrain evolution if within-cycle ecological dynamics reduce *N_e_*, or to differing interactions with *P. pacificus*. Further experiments should explore the ecology of this system in greater detail, including carrying capacity of the three organisms, as well as the ability of *P. pacificus* to prey on a diversity of *M. xanthus* genotypes to more fully characterize this ecological interaction and to identify *M. xanthus* genes and phenotypes relevant for anti-predator defense.

The evolutionary effects of an antibiotic-resistance mutation we observed raise intriguing questions for microbial mesopredators about possible evolutionary interactions between resisting higher-order predators, resisting antibiotic-mediated antagonisms from both conspecific and heterospecific microbes, and themselves preying on other microbes. It may be that resistance to antibiotics which target transcription may more generally come at a cost to an organism’s ability to adapt to predators. Even such an aggressive (Vos and Velicer 2009; Rendueles et al. 2015; Rendueles and Velicer 2017) and probably well-defended microbe as *M. xanthus* (Findlay 2016; Mayrhofer et al. 2021) is likely to experience trade-offs as it deals with antagonism from competitors and predators. Future research could examine potential trade-offs between competition within a trophic level (*e.g.,* competition among strains or species) and selective pressure from higher trophic levels, as well as the role of antibiotics and resistance to them in these conflicts (Cornforth and Foster 2015).

### Ecological context and the evolution of aggregative multicellularity

Because *M. xanthus* undergoes development in the abiotic selective regime used in this experiment (CFcc agar), particular evolutionary changes in developmental traits may be adaptive *per se.* For example, populations that evolved in the presence of *P. pacificus* may produce more fruiting bodies because they evolved to form fruiting bodies faster in order to better survive predation. Substantially reduced production by populations evolving with *E. coli* alone might reflect relaxed selection for sporulation if *E. coli* increases the level of resources available to *M. xanthus* by converting carbon sources used less efficiently *M. xanthus* into prey biomass. In this scenario, the phenotypic degradation we observed in the ME treatment could be due to generic adaptation to a high-resource environment (Velicer *et al*. 1998), rather than selection imposed by interactions with the prey itself. Testing among hypotheses to explain specific patterns of developmental-trait evolution across our treatments is of interest for future work.

*P. pacificus* drives the evolution of more fruiting bodies but not increased spore production (Figs. 4 and 5). In fact, spore production appears to respond as an independent trait to the presence of prey alone, as in La Fortezza and Velicer (La Fortezza and Velicer 2021), where spore production and fruiting body morphology evolved in an uncorrelated manner. This seeming independence suggests that the aggregation step of multicellular development may have evolved separately from spore formation, possibly in response to predation pressures (La Fortezza et al. 2022*b*). This could be tested specifically by evolving a strain which is defective at fruiting body formation in pure culture (Velicer et al. 2000; Schaal et al. 2022) in the presence of the predator. It would also be interesting to see for how long the difference in fruiting body numbers between the M and MP populations evolved here would be retained if both were to evolve further in conditions without the predator. Morphological trait integration analysis indicates that evolution under conditions of greater biotic complexity may support integration of *M. xanthus* developmental traits. However, the analysis presented here is speculative and further investigation is needed, in particular into the roles of *N_e_* and developmental constraint in this process and a potential genetic basis of trait integration.

Adaptation in predator-prey relationships spanning multiple trophic-level steps may be thought of as a process of escalation rather than one of mutual co-evolution (Vermeij 1987, 1994; Brodie and Brodie 1999). That is, changes in the morphology or behavior of a mesopredator (even for traits not directly related to predation) would drive responsive adaptation in its prey, but adaptation of the mesopredator would be driven largely by its own predators and not by the prey (Chattopadhyay and Dutta 2013; Klompmaker et al. 2017). A previous study has shown that prey identity can strongly affect how *M. xanthus* fruiting body phenotypes evolve indirectly (La Fortezza et al. 2022*a*) but in that study the prey did not co-evolve with *M. xanthus* (as also in this study), and fruiting body formation was not under direct selection. A subsequent evolution experiment would be of interest to test whether prey presence and type affect *mesopredator* AM evolution when an apex predator is present and when all parties can freely co-evolve, as well as to examine relative effects of the apex predator and basal prey.

How predation and multicellular development relate to each other in the evolutionary history of the myxobacteria is not yet well understood. Future comprehensive identification and phylogenetic analysis of the genes involved in both behaviors will improve understanding of their relative origins. While many selective forces are likely shape the evolution and diversification of aggregative multicellular systems (Bonner 1998; La Fortezza et al. 2022*b*), our results point to not only predation by mesopredators, but also predation on them, playing important roles.

## Authorship statement

KAS and GJV designed the evolution experiment. KAS, MLF, and GJV designed the fruiting body morphological analysis. KAS carried out the experimental work, except for the time lapse experiment. KAS, MLF, LMY, and GJV designed the time lapse experiment; LMY carried it out; LMY and MLF analyzed it. KAS performed the remaining statistical analysis. MLF advised on the statistical analysis. KAS, MLF, and GJV wrote the paper. KAS, MLF, LMY, and GJV revised the paper.

## Acknowledgements

We thank Hinrich Schulenburg, Ralf Sommer, and Peter Zee for advice and discussion. Some strains were provided by the CGC, which is funded by NIH Office of Research Infrastructure Programs (P40 OD010440). This research was supported in part by an EMBO Long- Term Fellowship (ALTF 1208–2017) to M.L.F. and in part by Swiss National Science Foundation (SNSF) grants 31003A_16005 and 310030B_182830 to G.J.V.

