## Supplemental Information for "Predation, evo-devo, and historical contingency: A nematode predator drives evolution of aggregative multicellularity"

Contents:

Figure S1. Eigenvector analysis of pleiotropy among developmental traits.

Figure S2. Spore production by fruiting body number and size.

Table S1. Two-way ANOVA results for linear correlation between spore counts and fruiting body number or size.

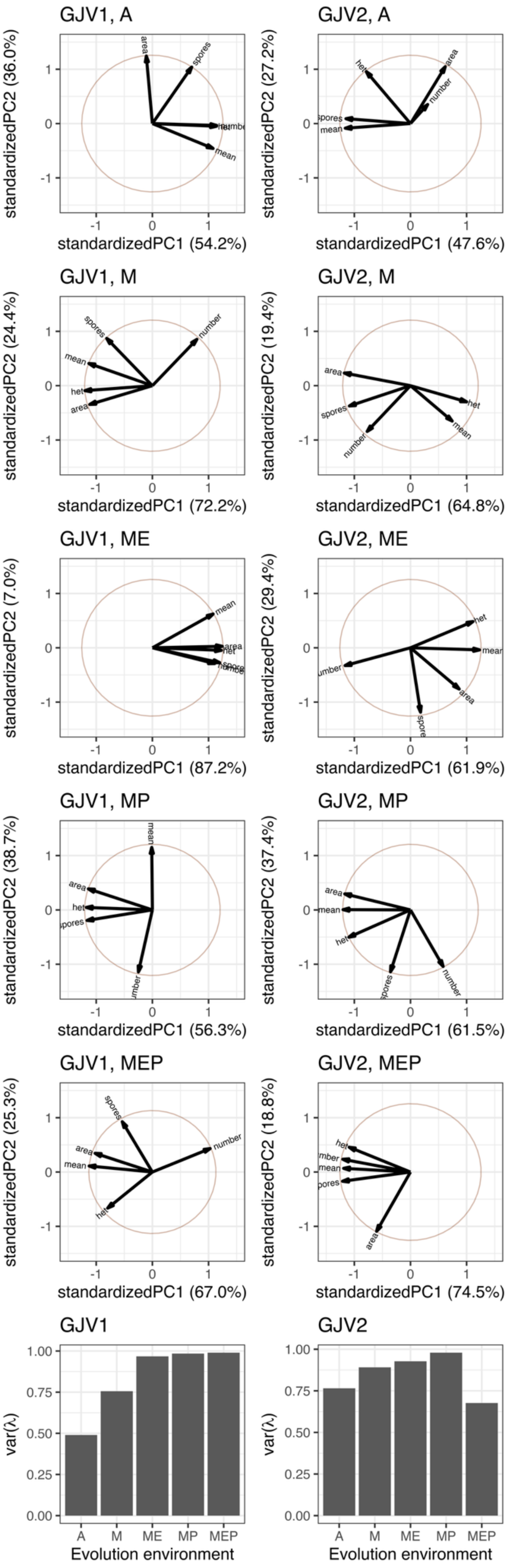

GJV1, A
standardizedPC1 (54.2%)
standardizedPC2 (36.0%)
area
spores
mean
GJV2, A
standardizedPC1 (47.6%)
standardizedPC2 (27.2%)
het
area
number
spores
mean
GJV1, M
standardizedPC1 (72.2%)
standardizedPC2 (24.4%)
spores
number
mean
het
area
GJV2, M
standardizedPC1 (64.8%)
standardizedPC2 (19.4%)
area
spores
number
het
mean
GJV1, ME
standardizedPC1 (87.2%)
standardizedPC2 (7.0%)
mean
area
GJV2, ME
standardizedPC1 (61.9%)
standardizedPC2 (29.4%)
het
mean
area
GJV1, MP
standardizedPC1 (56.3%)
standardizedPC2 (38.7%)
mean
area
het
GJV2, MP
standardizedPC1 (61.5%)
standardizedPC2 (37.4%)
area
mean
het
number
GJV1, MEP
standardizedPC1 (67.0%)
standardizedPC2 (25.3%)
spores
area
mean
number
het
GJV2, MEP
standardizedPC1 (74.5%)
standardizedPC2 (18.8%)
het
mean
area
GJV1
GJV2
var(λ)
Evolution environment
A
M
ME
MP
MEP

**Figure S1. Eigenvector analysis of pleiotropy among developmental traits.** Trait morphospaces of ancestral or evolved strains descending from GJV1 or GJV2, including fruiting body number, size, darkness, heterogeneity, and spore count; orthogonal arrows represent non-correlated traits. Bottom two panels show average eigenvector variance for the ancestral clones and each evolution environment. Traits: “het” = fruiting body heterogeneity, “area” = fruiting body size, “mean” = fruiting body darkness, “number” = number of fruiting bodies produced by a defined population of cells.

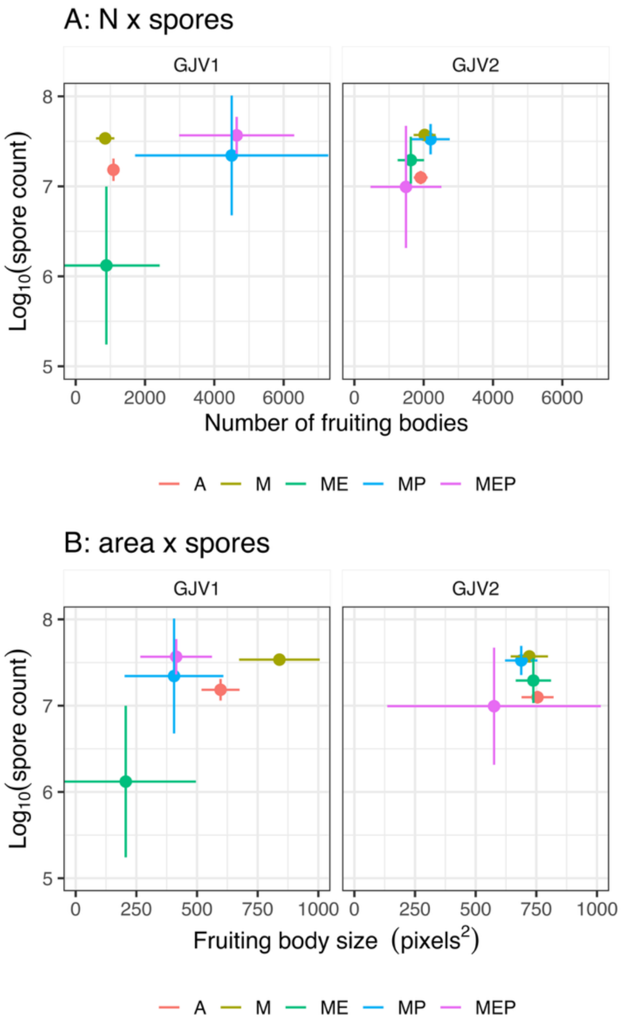


**Figure S2. Spore production by fruiting body number and size.** We show the relationship between spore production the number and size of fruiting bodies produced by descendants of GJV1 and GJV2. Dots represent the mean across 4-6 replicate lineages (each of which is calculated as a mean across 3-4 biological replicates), and bars show 95% confidence intervals of cross-lineage means. N = number of fruiting bodies, area = fruiting body size averaged within each plate, spores = number of spores produced. Spore counts are paired with the N or area value for the plate from which the spores were harvested. Panels group clones or populations descending from GJV1 or from GJV2.

**Table S1.** Two-way ANOVA results for linear correlation between spore counts and fruiting body number or size. df = degrees of freedom.

| Source | df | *F* | *p* |
|---|---|---|---|
| N CFcc GJV1 | | | |
| number | 1 | 21.97 | 0.0002 |
| treatment | 4 | 11.11 | 0.0002 |
| interaction | 4 | 2.05 | 0.1356 |
| residuals | 16 | | |
| N CFcc GJV2 | | | |
| number | 1 | 64.87 | < 0.0001 |
| treatment | 4 | 4.70 | 0.0090 |
| interaction | 4 | 2.52 | 0.0775 |
| residuals | 18 | | |
| area CFcc GJV1 | | | |
| area | 1 | 162.06 | < 0.0001 |
| treatment | 4 | 21.51 | < 0.0001 |
| interaction | 4 | 3.13 | 0.0442 |
| residuals | 16 | | |
| area CFcc GJV2 | | | |
| area | 1 | 9.06 | 0.0075 |
| treatment | 4 | 3.09 | 0.0424 |
| interaction | 4 | 0.19 | 0.9398 |
| residuals | 18 | | |
